# SIV/SHIV-Zika coinfection does not alter disease pathogenesis in adult non-pregnant Rhesus Macaques

**DOI:** 10.1101/347708

**Authors:** Mehdi R. M. Bidokhti, Debashis Dutta, Lepakshe S. V. Madduri, Shawna M. Woollard, Robert Norgren, Luis Giavedoni, Siddappa N. Byrareddy

## Abstract

Due to the large geographical overlap of populations exposed to Zika virus (ZIKV) and human immunodeficiency virus (HIV), understanding disease pathogenesis in such coinfections is urgently needed. We used chronically infected simian immunodeficiency virus and chimeric simian human immunodeficiency virus (SIV/SHIV) macaques and inoculated with ZIKV. Plasma viral loads of both SIV/SHIV and ZIKV showed no significant changes as compared to ZIKV alone-infected animals. Tissue clearance of ZIKV was observed similarly. Furthermore, minimal changes in cytokines/chemokines were observed. Collectively, these data suggest that coinfection may not alter disease pathogenesis and warrants large HIV-ZIKV epidemiological studies to validate these findings.

**Author Summary:** The co-infection incidence of human immunodeficiency virus (HIV) infection and neglected tropical infectious diseases is increasing due to the large geographical overlap of populations exposed to both of these viruses. Thus, researching on such coinfection is of particular importance. In this study, we investigated HIV-ZIKV coinfection dynamics in adult non-pregnant Rhesus Macaques model chronically infected with simian immunodeficiency virus (SIV) - or chimeric simian human immunodeficiency virus (SHIV). We found that post ZIKV inoculation, plasma viral loads were similar to ZIKV alone infected animals in addition to minimal changes of cytokines. Dynamics of SIV and SHIV also did not change. Tissue clearance of ZIKV was found 6-7 months later. Our findings provide insights into HIV-ZKIV coinfection to determine the alteration of their pathogenesis.

## Introduction

Experimental and theoretical attention has been paid to the interactions between human immunodeficiency virus (HIV) infection and neglected tropical infectious diseases. Such interactions are marked by the potential that a pathogen has for changing the epidemiology, pathogenesis, immunology, and response to therapy of the other **[1]**. In the last six decades since its discovery, Zika virus (ZIKV) has been considered a mild human pathogen; but recently, it has emerged as a threat to global health, showing increased virulence, rapid spread and an association with grave neurological complications **[2, 3]**. The two major clinical complications for ZIKV infection are microcephaly of newborns from women infected during early pregnancy **[4]**, and neurological conditions in adults, including Guillain-Barré syndrome **[1]**. Serological tests cross-react with dengue, and there are neither specific antivirals nor vaccines. The major tool to combat ZIKV is prevention against mosquito bites using repellents and insecticides **[1]**.

The association between HIV infection and endemic diseases has been described in tropical regions with different levels of complications. The first case of HIV-ZIKV coinfection was reported in Brazil without major health complications **[5]**. However, as the geographical range of ZIKV infection expands, exposed HIV immunosuppressed individuals may unveil new and more severe clinical manifestations. Therefore, close surveillance of HIV-positive individuals to mirror such coinfections is of particular importance **[1]**. We investigated SIV/SHIV-ZIKV coinfection dynamics in a biologically relevant nonhuman adult non-pregnant primate model to test whether the status of these two viral infections could be changed. Our aim was to determine if ZIKV infection in HIV positive individuals has any altered pathogenesis.

## Methods

### Animals and study design

As described in **Fig 1 A**, total of 6 adult female Indian-origin rhesus macaque (*Macaca mulatta;* age range 4.5 to 5 yrs) were chronically infected with SIVmac239 or SHIV3618MTF over a period of 6-7 months **(Table 1)**. These animals were inoculated subcutaneously with 10^4^ plaque forming unit (PFU) of ZIKV strain PRVABC59 **(Table 1)** and monitored for post ZIKV infection by viral loads and also evaluated for any clinical manifestation caused by ZIKV. All Animals studies were conducted in accordance with UNMC IACUC approved protocols. Animal maintenance and procedures were done at the Department of Comparative Medicine, University of Nebraska Medical Center (UNMC) in accordance with the rules and regulations of the Committee on the Care and Use of Laboratory Animal Resources, and according to the guidelines of the Committee on the Care and Use of Laboratory. All protocols and procedures were performed under approval of the UNMC Institutional Animal Care and Use Committee according to the National Institute of Health guidelines.

**Fig 1.**
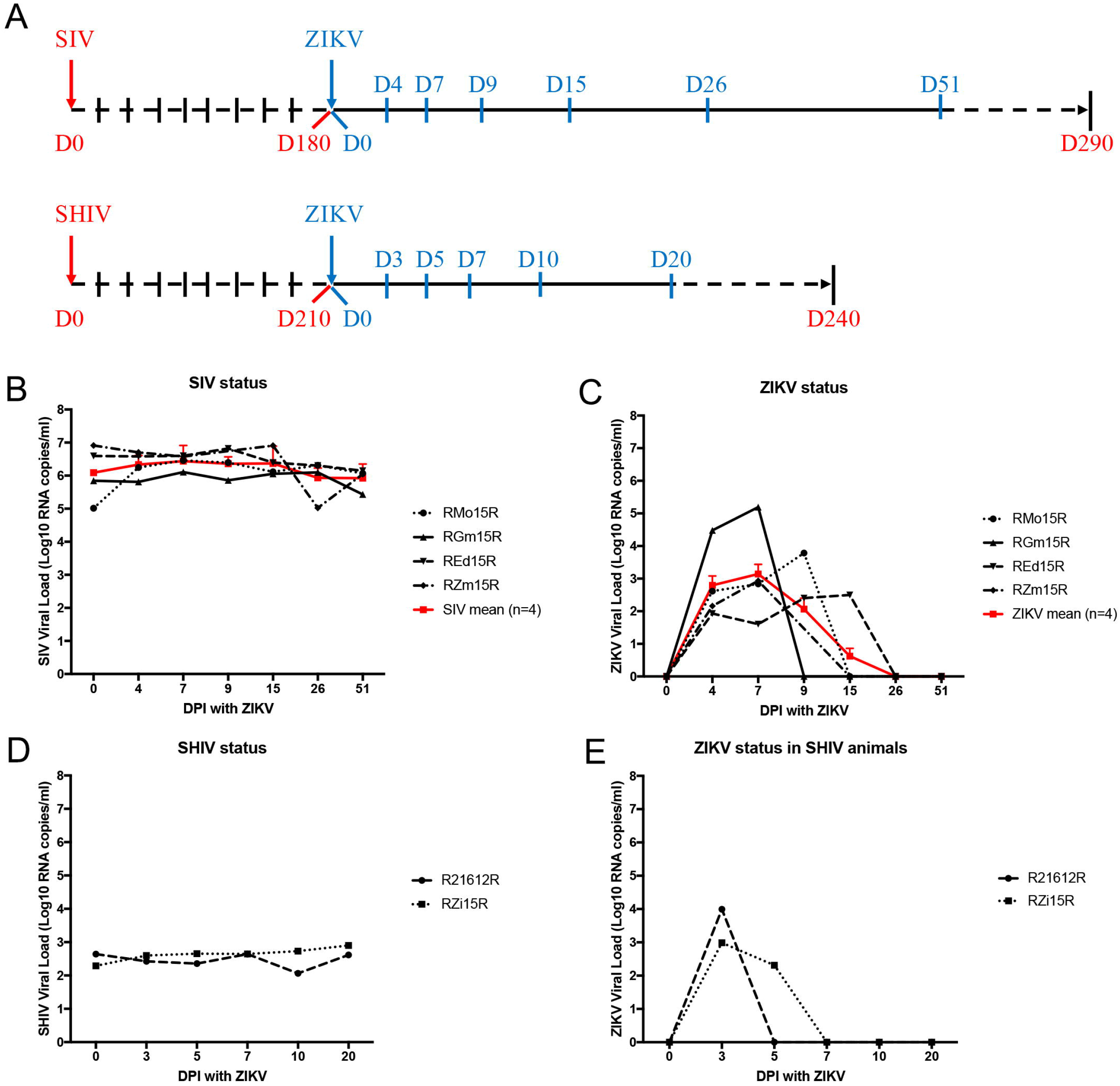
Viral loads of simian immunodeficiency virus and chimeric simian human immunodeficiency virus (SIV/SHIV) and Zika virus (ZIKV) in co-infected monkeys. Rhesus macaques (n = 6) chronically infected with SIVmac239 (n = 4) or SHIV3618MTF (n = 2) were also inoculated subcutaneously with 10^4^ plaque forming unit (PFU) of ZIKV PRVABC59. Blood collection was performed according to the study plan on 0, 4, 7, 9, 15, 26 and 51 days post inoculation (dpi) with ZIKV for SIV coinfected individuals and on 0, 3, 5, 7, 10 and 20 DPI with ZIKV for SHIV co-infected individuals. Day 0 (D0) was the day of inoculation with ZIKV. From collected plasma samples, RNA was extracted using QiAmp RNA mini kit (Qiagen, Valencia, CA), and viral loads were measured using one-step real time RT-PCR detection method targeting Gag gene of SIV. Viral loads were presented in Log10 RNA copies per milliliter (ml) of plasma. **A**, schema of time-course sampling in study plan of SIV (n = 4) and SHIV (n = 2) coinfection with ZIKV in rhesus macaques. **B**, viral load status of SIV in individual (black) and the mean value (red) of all SIV-ZIKV co-infected four animals (n = 4) in days post ZIKV inoculation. **C**, viral load status of ZIKV in individual (black) and the mean value (blue) of all SIV-ZIKV co-infected animals (n = 4) in days post ZIKV inoculation. Bars indicate standard deviation (±SD) of mean values (n=4). **D**, viral load status of SHIV in individual SHIV-ZIKV co-infected animals (n = 2) in days post ZIKV inoculation. **E**, viral load status of ZIKV in individual SHIV-ZIKV co-infected animals (n = 2) in days post ZIKV inoculation.

**Table 1.** Information about monkey animals used in this study.

| Name of animals | Species of rhesus macaques | Type of inoculated immuno-deficiency virus isolate | Type of inoculated ZIKV isolate | Gender | Date of birth | Date and route of SIV or SHIV inoculation | Weight on date of SIV or SHIV infection | Date and route of ZIKV inoculation | Date of Necropsy |
| --- | --- | --- | --- | --- | --- | --- | --- | --- | --- |
| Mo15R | Mulatta | SIVmac239 | ZIKVPRV ABC59 | Female | 6/25/2012 | 3/29/2016, IV<sup>a</sup> | 5.54kg | 9/26/2016, SC<sup>c</sup> | 1/12/2017 |
| Gm15R | Mulatta | SIVmac239 | ZIKVPRV ABC59 | Female | 5/30/2012 | 3/29/2016, IV | 5.52kg | 9/26/2016, SC | 1/12/2017 |
| Ed15R | Mulatta | SIVmac239 | ZIKVPRV ABC59 | Female | 6/12/2012 | 3/29/2016, IV | 6.12kg | 9/26/2016, SC | 1/12/2017 |
| Zm15R | Mulatta | SIVmac239 | ZIKVPRV ABC59 | Female | 6/7/2012 | 3/29/2016, IV | 5.46kg | 9/26/2016, SC | 1/17/2017 |
| 21612R | Mulatta | SHIV3618M TF | ZIKVPRV ABC59 | Female | 7/30/2012 | 9/20/2016, IVAG<sup>b</sup> | 5.54kg | 4/21/2017, SC | 5/11/2017 |
| Zi15R | Mulatta | SHIV3618M TF | ZIKVPRV ABC59 | Female | 5/11/2012 | 9/20/2016, IVAG | 5.56kg | 4/21/2017, SC | 5/10/2017 |

<sup>a</sup> Intravenously.114 <sup>b</sup> Intravaginally.115 <sup>c</sup> Subcutaneously.

### Sample collection and viral loads in plasma and tissues

On various days post inoculation (dpi) with ZIKV **(Fig 1A)**, blood samples were collected and centrifuged to obtain plasma. At necropsy 6-7 months post ZIKV infection, after perfusion of blood, organs were harvested and snap frozen for RNA extraction. From collected plasma samples, RNA was extracted using QiAmp Viral RNA Mini Kit (Qiagen, CA). Quantitation of viral RNA was performed using the Taqman^®^ RNA-to-Ct^TM^ 1-Step Kit (Thermo Fisher Scientific, MA), and SIV and ZIKV viral loads were measured as described previously **[6, 7]**. Standard curves of SIV and ZIKV quantitative real-time PCR are presented in **S1A and S1B Figs**, respectively. Similarly, to measure ZIKV, RNA was extracted from plasma/various tissues and subjected to sensitive QX200 Droplet Digital PCR assay with limit of detection as low as 3 copies/ml (Bio-Rad, CA). A reaction mixture containing 200 ng of DNA was made using the One-Step RT-ddPCR Advanced Kit for Probes (Bio-Rad, Hercules CA). Microdroplets were generated using the QX200 Automated Droplet Generator (Bio-Rad, Hercules CA). Plates were sealed with the PX1 PCR Plate Sealer (Bio-Rad, Hercules CA) prior to PCR. Target DNA was amplified with the C1000 Touch Thermal Cycler (Bio-Rad, Hercules CA) using the following conditions: 1 cycle 48°C for 1 hour, 1 cycle 95°C for 10 min, 40 cycles 95°C for 30 secs and 60°C for 1 min, 1 cycle 98°C for 10 min. After amplification, the plate was read on a QX200 Droplet Reader (Bio-Rad, Hercules CA) to determine the number of PCR-positive droplets vs PCR-negative droplets in the original sample. Data acquisition and quantification was performed using QuantaSoft Software (Bio-Rad, Hercules CA). Furthermore, ZIKV viral loads from plasma of SIV-ZIKV confected animals were compared with those of the ZIKV alone infected animals from previous study **[8]**.

### Levels of cytokines and chemokines

Plasma samples from SIV-ZIKV co-infected macaques were analyzed, by commercially available Luminex methodology **[9]**, for measuring the levels of cytokines and chemokines including: B lymphocyte chemoattractant (BLC)/CXCL13, Eotaxin, interferon-γ-induced protein 10 (IP-10), monocyte chemoattractant protein-1 (MCP-1), regulated on activation, normal T cell expressed and secreted (RANTES), interferon-inducible T-cell alpha chemoattractant (I-TAC), macrophage migration inhibitory factor (MIF), interferon (IFN)-α, IFN-γ, interleukin (IL)-1β, IL-6,, IL-7, IL-8, stromal cell-derived factor (SDF)-1α, IL-1Rα, and GRO-α, **(Fig 2**, **S1 Table)**.

**Fig 2.**
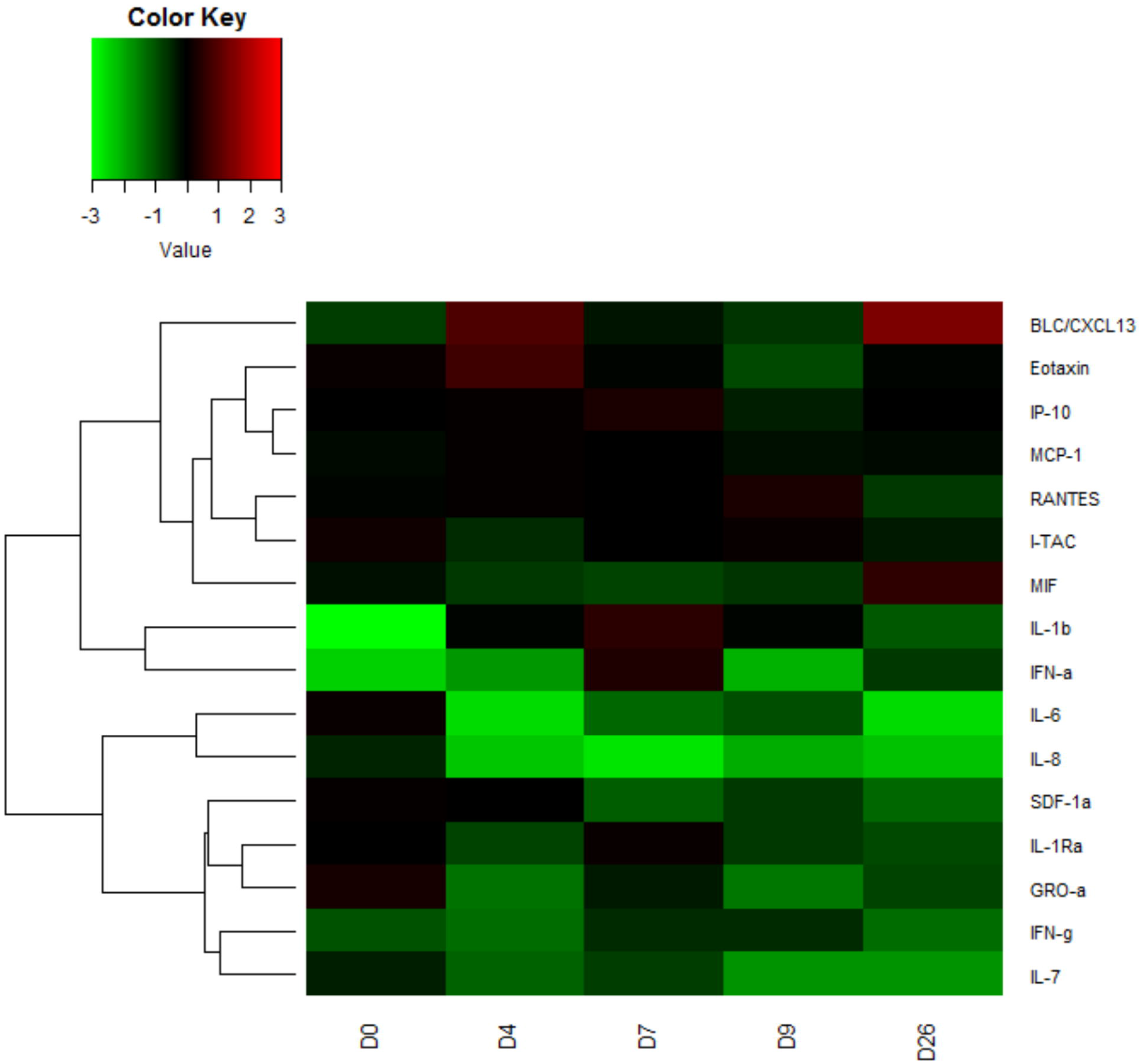
The heatmap of cytokines/chemokines measurement screening in simian immunodeficiency virus - Zika virus (SIV-ZIKV) co-infected monkeys. Rhesus macaques (n = 3) chronically infected with SIVmac239 were also inoculated with 10^4^ plaque forming unit (PFU) of ZIKV PRVABC59 subcutaneously. The blood collection was performed according to the study plan on 0, 4, 7, 9, 15, 26 and 51 days post inoculation (dpi) with ZIKV. Day 0 (D0) was the day of inoculation with ZIKV. From collected serum samples on D0 to D26 post inoculation with ZIKV, Luminex assay was performed to screen all relevant cytokines/chemokines measurements. Using R, the heatmap was designed and drawn based on the median value (n = 3) of cytokines/chemokines measurement on D0, D4, D7, D9, and D26 post inoculation with ZIKV in SIV-ZIKV co-infected animals. Raw values for each of the cytokines/chemokines were normalized to the mean of the baselines (**S1A Table**), log2-transformed (**S1B Table**) and then subjected to hierarchical clustering; heat maps were generated using heatmap.2 of R package plots. Values denoted as −3 to +3 represent a decrease to an increase in the levels relative to base line values for each of chemokine/cytokine analyzed. BLC, B lymphocyte chemoattractant; IP-10, interferon-β-induced protein 10; MCP-1, monocyte chemoattractant protein-1; RANTES, regulated on activation, normal T cell expressed and secreted; I-TAC, interferon-inducible T-cell alpha chemoattractant; MIF, macrophage migration inhibitory factor; IFN--α and −γ, interferon-α and −γ; IL, interleukin; SDF-1α, stromal cell-derived factor.

### Statistical analysis

ANNOVA single, two-way without replication and two-way with replication were used to compare various data presented in this study.

## Results

### SIV/SHIV and Zika plasma/tissue viral load measurements

First, plasma viral loads of all the chronically infected SIV macaques were stable (10^5^-10^7^ copies/ml) and maintained as before ZIKV inoculation as shown in SIVmac239 **(Fig 1B**). However, their ZIKV plasma viral loads were found to peak up to 10^5^ copies/ml on 7 dpi **(Fig 1C**). ZIKV was detected in plasma samples up to 9 dpi in two of SIV-ZIKV co-infected animals; <10^4^ copies/ml for RMo15R and > 10^2^ copies/ml for REd15R on 9 dpi. RGm15R that had the highest viral load, > 10^5^ copies/ml on 7 dpi, and was found negative to ZIKV on 9 dpi **(Fig 1C**). Interestingly, REd15R, with the lowest peak of ZIKV viral load of > 10^2^ copies/ml, detected ZIKV up to 15 dpi **(Fig 1C**). Furthermore, the SIV viral load status of these four SIV-ZIKV co-infected RMs were similar and quite stable during 51 dpi with ZIKV **(Fig 1B**).

In SHIV-ZIKV confected RMs, SHIV plasma viral loads were also found quite stable 10^2^-10^3^ copies/ml during 20 dpi with ZIKV **(Fig 1D**). Also, their ZIKV plasma viral loads were found to peak on 3 dpi to 10^3^ copies/ml for R21612R animal and 10^4^ copies/ml for RZi15R animal **(Fig 1E**). On 5 dpi, ZIKV was only detected at lower levels, < 10^3^ copies/ml, in RZi15R; after that and up to end of study, ZIKV was not found in RMs R21612R and RZi15R **(Fig 1E**).

Viral loads of ZIKV in all six co-infected RMs in this study were found negative after 20 days onwards; however, this delay of self-recovery of viremia is longer in coinfected animals as compared to ZIKV alone-infected animals (**S2 and S3 Figs)**. For statistical analysis, ANNOVA single (p value=0.48), two-way without replication (p value=0.42) and two-way with replication (p value=0.51) of viral load of ZIKV did not show significant difference between co-infected and alone-infected RMs inoculated with 10^4^ PFU of ZIKV PRVABC59. Lower number of studied animals does not provide enough evidence to confirm this delay of self-recovery of ZIKV viremia (**S2B Fig**), which requires more epidemiological studies. Clinical investigation of co-infected RMs did not show any severe symptoms and/or clinical signs of ZIKV infection with permanent sequelae after **acute phase (≤ 9 dpi)**. Additionally, at necropsy 6-7 months post inoculation with ZIKV, various tissues including brain stem, hippocampus, caudate, cerebellum, frontal cortex, spleen, mesenteric lymph node, uterus, liver, lung, and kidney, were collected and tested for ZIKV detection using a highly sensitive Droplet Digital PCR. We found that no detectable ZIKV virus was present in any of the sampled tissues of three SIV-ZIKV co-infected RMs (RMo15R, RGm15R, and REd15R) suggesting that ZIKV infection had been cleared in these animals **(data not shown)**. This finding indicates the clearance of the ZIKV infection from the SIV-ZIKV co-infected adult RMs as similar to previously reported studies for ZIKV alone **[10, 11]**.

### Measurement of cytokines/chemokines

Using Luminex methodology **[9]**, cytokines and chemokines were measured in plasma samples of three SIV-ZIKV co-infected RMs (RMo15R, RGm15R, and REd15R) on 0, 4, 7, 9, and 26 dpi with ZIKV. The results show that with the exception of MIF, IL-8, and SDF-1α, significant elevations of the serum interleukin concentrations are evident in the acute phase of ZIKV especially at the peak on 7 dpi. Many of the cytokines and chemokines that were elevated in the acute phase of ZIKV showed a tendency to return to normal levels in the later **recovery phase (26 dpi)** of ZIKV. BLC/CXCL13, Eotaxin, IP-10, MCP-1, and IFN-a were elevated during the peak of viremia (D4-7) and also in recovery phase (D26). RANTES, I-TAC and, at lower levels, IL-1β, IL-6, IL-7, IFN-γ, SDF-1α, IL-1Rα, and GRO-α were elevated during acute phase and suppressed in recovery phase. Elevation of IL-1β and IFN-α in acute phase was noted. MIF and, at lower levels, IL-8 were suppressed during acute phase and elevated in recovery phase. The changes, either in the acute or in the recovery phase, were minor for MCP-1, IL-5, and IL-7 **(Fig 2**, **S1 Table)**.

## Discussion

The main objective of this study is to understand the dynamics of HIV-ZIKV coinfection and whether one pathogen changes course of the disease and pathogenesis of the other. We used rhesus macaques chronically infected with either SIVmac239 or SHIV3618MTF. These macaques were then inoculated with 10^4^ PFU of ZIKV (PRVABC59) subcutaneously. In SIV-ZIKV co-infected RMs, Zika viral loads in plasma were found to be very similar to ZIKV infected animals in the literature **[8]**, as well as our own data (unpublished). Plasma viral loads of SIV and SHIV did not change as compared prior to ZIKV inoculation. These levels of viral load status of SIV and SHIV have been observed in our previous studies as well as chronically infected RMs in literature. **[12, 13]**. Generally, mosquito-borne flaviviruses are initially detected in blood and lymphoid tissue of infected animals and subsequently appear in peripheral organs and invade central nervous system via the hematogenous route **[1, 2, 8]**. In chronically infected SIV/SHIV RMs, ZIKV (PRVABC59) exhibited a similar pattern of viral kinetics as previously described in the ZIKV infected animals **[8, 11]**. The SIV/SHIV viremia kinetics in these co-infected RMs were also similar to those in the SIV/SHIV infected RMs **[12, 13]**. This rapid control of acute viremia of ZIKV infection in SIV/SHIV chronically infected RMs suggests that their peripheral immune system protects the host from peripheral ZIKV infection and that chronically infected SIV/SHIV RMs are similarly protected as non-immunocompromised RMs from infection by ZIKV.

Next, necropsy was performed 6-7 months post ZIKV infection, and Zika viral loads were measured in different tissues and organs using a highly sensitive Droplet Digital PCR with no ZIKV detection results. Previous studies also showed similar clearance from tissue organs and body fluids for ZIKV (PRVABC59) infection in human **[10]** and in RMs **[11]**. Human and animal model studies have demonstrated that ZIKV infection can result in persistence of infectious virus and viral nucleic acid in several body fluids (e.g., semen, saliva, tears, and urine) and target organs, including immune-privileged sites (e.g., eyes, brain, and testes) and the female genital tract **[11, 14]**. The ZIKV persistent or occult neurologic and lymphoid disease may occur following clearance of peripheral virus in ZIKV-infected individuals **[11]**. It has been shown in infected RMs that ZIKV can persist in cerebrospinal fluid and lymph nodes for weeks after virus has been cleared from peripheral blood, urine, and mucosal secretions **[11]**. The present adult RMs model confirmed that ZIKV persistent infection would also be cleared in immunocompromised RMs chronically infected with SIV. However, we didn’t investigate the rate of ZIKV clearance if it happened earlier than 6-7 months. ZIKV infection of rhesus and cynomolgus monkeys has been shown to recapitulate many key clinical findings, including rapid control of acute viremia, early invasion of the central nervous system, prolonged viral shedding in adult animals **[8, 11, 14]**. In the present study, rapid control of acute viremia of ZIKV infection in SIV/SHIV chronically infected RMs was also detected. Collectively, these findings enumerate that ZIKV infection in HIV patients and immunocompromised individuals would not alter significant changes in pathogenesis and treatment plans.

Furthermore, except MIF, IL-8, and SDF-1α, the cytokine/chemokine patterns were elevated during acute phase of ZIKV infection in RMs. Interestingly, this pattern with minimal changes is in accordance with previously described clinical findings for ZIKV-infected RMs **[13]** and human patients **[15]**. Major elevation of IL-1β and IFN-α in acute phase of ZIKV infection was observed in SIV-ZIKV co-infected RMs. The cytokine IL-1β is a key mediator of the inflammatory response and is essential for the host-response and resistance to pathogens. IFN-α is also mainly involved in innate immune responses against viral infection in acute phase of infection **[9]**. Collectively, these data confirm that ZIKV replication in the acute phase triggered rapid innate immune responses in peripheral blood. Several chemoattractant chemokines (BLC/CXCL13, IP-10, MCP-1, RANTES, I-TAC, SDF-1α, and GRO-α) were also elevated during the ZIKV infection in chronically SIV/SHIV infected RMs described in this study. These chemokines induce protective immunity against various viral infections including influenza, herpes simplex virus, Coxsackie virus, respiratory syncytial virus, and flaviviruses such as dengue virus and west Nile virus **[1, 15]**. Elucidating the function of these chemokines in ZIKV infection, such as trafficking of lymphocytes into the various tissues, may explain mechanisms of immunological protection or immunopathology in SIV/SHIV-ZIKV coinfection. For rapid control of infection, both B cell and T cell recruitment to the sites of infection were highly triggered in SIV-ZIKV co-infected RMs. However, there were no significant changes between any of the cytokine levels measured in the acute versus the recovery phase.

In summary, we showed that ZIKV viremia dynamics in chronically SIV/SHIV RMs did not change much when compared to ZIKV alone infected RMs and human patients. While these data suggest that ZIKV infection in chronically infected HIV individuals may not significantly alter the pathogenesis and disease progression of HIV or ZIKV, this study warrants more epidemiological studies to validate these findings. The results help with better planning vaccination and treatment strategies.

## Supporting information

**S1 Fig. Standard curve of quantitative real-time PCR.** In this study to quantify viral loads of SIV/SHIV and ZIKV in plasma samples collected in various time point samplings, quantitative real-time PCR assays were used. **A)** Standard curve of real-time PCR to quantify SIV viral loads. **B)** Standard curve of real-time PCR to quantify ZIKV viral loads.

**S2 Fig. Comparison of viral loads in simian immunodeficiency virus and chimeric simian human immunodeficiency virus - Zika virus (SIV/SHIV-ZIKV) co-infected monkeys versus only ZIKV-infected monkeys.** Rhesus macaques (RM; n = 6) chronically infected with SIVmac239 (n = 4) or SHIV3618MTF (n = 2) were also inoculated subcutaneously with 10^4^ plaque forming unit (PFU) of ZIKV PRVABC59. The blood collections were performed according to the study plan on 0, 4, 7, 9, 15, 26 and 51 days post inoculation (DPI) with ZIKV for SIV co-infected RM and on 0, 3, 5, 7, 10, and 20 DPI with ZIKV for SHIV co-infected RM. Day 0 (D0) was the day of inoculation with ZIKV. From collected plasma samples, RNA were extracted using QiAmp RNA mini kit (Qiagen, Valencia, CA), and viral loads were measured using one-step real time RT-PCR detection method targeting Gag gene of SIV. Viral loads were presented in Log10 RNA copies per milliliter (ml) of plasma. **A)** Viral load status of ZIKV in SIV (black, this study), SHIV (orange, this study) ‒ ZIKV co-infected RM, in ZIKV PRVABC59 (red) - infected RM, in ZIKV FrenchPolynesia,2013 (brown) - infected RM and in ZIKV MR766 (blue) - infected RM in days post ZIKV inoculation. The reference number for animals studied previously are also given in their labels (https://zika.labkey.com/project/OConnor/ZIKV); 295022 (infected with 10^4^ PFU of ZIKV MR766); 912116 and 411359 (both infected with 10^4^ PFU of ZIKV French Polynesia isolated in 2013); 634675, 566628, and 311413 (all infected with 10^4^ PFU of ZIKV PRVABC59 isolated in Puerto Rico in 2015). **B)** Mean value of viral load status of ZIKV (strain PRVABC59) in all SIV-ZIKV co-infected animals (n = 4, blue) and in all ZIKV PRVABC59 infected animals (n = 3, red) in days post ZIKV inoculation. Bars indicate standard deviation (± SD) of mean values.

**S3 Fig. Body weight and temperature status of simian immunodeficiency virus and chimeric simian human immunodeficiency virus - Zika virus (SIV/SHIV-ZIKV) coinfected monkeys versus ZIKV-infected monkeys (from David O’Connor NIH research group).** Rhesus macaques (n = 6) chronically infected with SIVmac239 (n = 4) or SHIV3618MTF (n = 2) were also inoculated subcutaneously with 10^4^ plaque forming unit (PFU) of ZIKV PRVABC59. The blood collections, body weight, and temperature records were performed according to the study plan on 0, 4, 7, 9, 15, 26 and 51 days post inoculation (DPI) with ZIKV for SIV co-infected animals and on 0, 3, 5, 7, 10 and 20 DPI with ZIKV for SHIV co-infected individuals. Day 0 (D0) was the day of inoculation with ZIKV. **A)** Body weight (Kg) records of individual animals on days pre and post inoculation with ZIKV. The reference number of the animals studied previously are also given in their labels (https://zika.labkey.com/project/OConnor/ZIKV); 295022 (infected with 10^4^ PFU of ZIKV MR766); 912116 and 411359 (both infected with 10^4^ PFU of ZIKV French Polynesia isolated in 2013); 634675, 566628, and 311413 (all infected with 10^4^ PFU of ZIKV PRVABC59 isolated in Puerto Rico in 2015). **B)** Mean value of body weight (Kg) records of all SIV-ZIKV co-infected animals (n = 4, blue) and in all ZIKV PRVABC59 infected animals (n = 3, red) in days pre-and post ZIKV inoculation. Bars indicate standard deviation (±SD) of mean values. **C)** Body temperature (Celsius) records of individual animals on days pre-and post-inoculation with ZIKV. The reference number of the animals studied previously are also given in their labels (https://zika.labkey.com/project/OConnor/ZIKV); 295022 (infected with 10^4^ PFU of ZIKV MR766); 912116 and 411359 (both infected with 10^4^ PFU of ZIKV French Polynesia isolated in 2013); 634675, 566628, and 311413 (all infected with 10^4^ PFU of ZIKV PRVABC59 isolated in Puerto Rico in 2015). **D)** Mean value of body temperature (Celsius) records of all SIV-ZIKV co-infected animals (n = 4, blue) and in all ZIKV PRVABC59 infected animals (n = 3, red) in days pre-and post ZIKV inoculation. Bars indicate standard deviation (±SD) of mean values.

**S1 Table. Raw data of cytokines/chemokines measurement screening in simian immunodeficiency virus - Zika virus (SIV-ZIKV) coinfected monkeys.** Rhesus macaques (n = 3) chronically infected with SIVmac239 were also inoculated subcutaneously with 10^4^ plaque forming unit (PFU) of ZIKV PRVABC59. The blood collections were performed according to the study plan on 0, 4, 7, 9, 15, 26 and 51 days post inoculation (DPI) with ZIKV. Day 0 (D0) was the day of inoculation with ZIKV. From collected serum samples on D0 to D26 post inoculation with ZIKV, Luminex assay was performed to screen all relevant cytokines/chemokines measurements. Using R, the heatmap was designed and drawn based on the median value (n = 3) of cytokines/chemokines measurements on D0, D4, D7, D9 and D26 post inoculation with ZIKV in SIV-ZIKV confected animals. **A)** Raw data of Luminex assay. **B)** Log2 of the raw data of which median values were calculated and used to draw the heatmap **(Fig 2)** using R program software.

